# Highly efficient healing of critical sized articular cartilage defect in situ using a chemically nucleoside-modified mRNA-enhanced cell therapy

**DOI:** 10.1101/2022.05.06.490932

**Authors:** Gang Zhong, Yixuan Luo, Jianping Zhao, Meng Wang, Fan Yang, Jian Huang, Lijin Zou, Xuenong Zou, Qingqing Wang, Fei Chen, Gang Wang, Yin Yu

## Abstract

Critical sized cartilage defects heal poorly and MSC-based therapies holds promise functional cartilage regeneration either used alone or in combination with growth factors. However, Recombinant protein growth factors were proven to have minimal benefits while to have adverse side effects and high cost. Nonviral mRNA delivery provides a promising, alternative approach to delivering therapeutic proteins within defect lesion for an extended period of time. Despite successful therapeutic outcome in bone and other vascularized tissues, the therapeutic application of mRNA in poorly vascularized tissues such as cartilage is still facing many challenges and rarely studied. We report here using chemically modified messenger RNA encoding TGF-β3(TGF-β3 cmRNA) to enhance the therapeutic efficacy of BMSCs to efficient repair of cartilage defect. Local administration of TGF-β3 cmRNA enhanced BMSCs therapy restored critical-sized cartilage defects in situ in a rat model within 6 weeks with structural and molecular markers similar to its nature counterparts. In addition, the development of osteoarthritis caused by cartilage damage was prevented by this mRNA-enhanced BMSCs therapy evidenced by minimal late-stage OA pharmacological presentations. This novel mRNA enhanced-MSC technology extend the development of new therapeutic approaches for treating functional cartilage repair.

## Introduction

Articular cartilage is an important component of synovial joint, cushioning the vibrations and shocks of connected bones during walking, jumping, and other movements[1], which makes it prone to mechanical injury in rigorous athletic and recreational activities[2, 3]. As an avascular tissue, which lacks blood and nutrient supply, articular cartilage once damaged can hardly regenerate by itself, which eventually leads to osteoarthritis (OA)[4], which is the most common chronic joint ailments worldwide, with estimates showing that it will affect 78 millions people by 2040[5, 6].

High incidence and limited self-healing pose challenges for the treatment of cartilage defects in both young atheletic population and elderly individuals. Traditional treatments, including nonsteroidal drugs, physiotherapy and surgical transplantation (autograft or allograft), are hindered by their shortcomings such as inefficiency, long-term treatment, side effects and immune rejection[7, 8]. Stem cell therapy has recently received extensive attention for musculoskeletal regeneration due to its multilineage differentiation potential which has demonstrated enormous tissue repair effects in numerous attempts[9].

However, the multilineage potential of stem cells is a double-edged sword, conferring the therapeutic efficacy of stem cell therapy in a variety of tissues, while retaining the risk of teratoma and carcinogenesis, which brings huge potential safety concerns to stem cell therapy[10, 11]. Several researchers have described the role of MSCs transplantation in tumor formation. Tolar and colleagues transplanted MSCs into nude mice and observed tumor formation, and the formed tumors exhibited characteristics typical of osteosarcoma, including sarcoma morphology, typical osteoid formation, and the prevalence of lung metastases[12]. Mohseny et al continuously cultured freshly obtained BMSCs in vitro and injected them into mice subcutaneously. As a result, osteosarcoma-like changes were observed in the back of the mice. Further mechanistic analysis revealed that the continuous loss of the Cdkn2 gene in MSCs appeared to be a key event in the carcinogenesis of MSCs during in vitro culture[13]. To sum up, the unguided and unfettered implementation of stem cell therapy may carry enormous risks and lead to serious consequences.

Controlled stem cells differentiation involves many aspects, especially growth factors, which serves as an outstanding candidate to guide stem cell therapy by mediating directed differentiation in a targeted lineage, and some of them have been utilized in clinic, showing great efficacy[14]. Despite the proven success, this therapeutic strategy was shelved in clinical scenarios, largely blaming its short half-life when contacted with body fluids and tissues[15, 16] as well as growing tendency of immune reactions overtime. Long term superphysiological dosage of protein growth factors have shown evident side effects[17].

Gene therapy provides a promising alternative to supply therapeutic proteins in a moderate yet sustainable manner. As an initial attempt with limitations, pDNA is integrated into the genome of target cells with the help of vectors to achieve continuous translation of therapeutic proteins[18]. However, the low transfection efficiencies due to the need for nuclear entry and safety concerns are potential barriers for their clinical translation[19]. Messenger RNA (mRNA) has recently been shown to be an attractive alternative to pDNA. Unlike the case with DNA therapeutics, its effector site is in the cytoplasm and the encoded therapeutic protein is immediately translated in the ribosome with no need to translocate to the nucleus of the cell, which significantly decreases the risk of insertional mutagenesis[20]. Remarkably, mRNA might be capable of enhancing cell therapy. For instance, a single leukapheresis derived peripheral blood lymphocytes are transfected with anti human mesothelin mRNA CAR and are stored as multiple cell aliquots for repeat transfusion[21]. Currently, mRNA enhanced cell therapy(CAR-T) have progressed into clinical trials[22]. Together, recent advances in mRNA technology, led by molecular backbone modifications and mRNA enhanced cell therapy such as the new CAT-T therapy[23-25] have sparked very strong interests in the use of mRNA in regenerative medicine as a novel, safe and effective way for tissue regeneration.

Given that transforming growth factor-β3 (TGF-β3) plays a key role in cartilage regeneration, capable of inducing chondrogenic differentiation of MSCs and promoting cartilage-like matrix deposition[26], it was selected as targeted protein of which gene encoded for in this project. Here, we have evaluated the ability of a chemically modified mRNA encoding TGF-β3 (TGF-β3 cmRNA) to repair the cartilage defects in a rat articular cartilage defect model by comparing with BMSCs-only treatment strategies. The data demonstrate TGF-β3 cmRNA-enhanced cell therapy effectively healed a critical-sized defect in the rat articular cartilage in situ. The work summarized here illustrates the efficacy and safety of an affordable mRNA-enhanced cell therapy for cartilage regeneration in situ and also hold implication for other low vascularized tissue repair.

## Materials and methods

### Isolation and culture of rat BMSCs

Briefly, Bone marrow was collected from the femoral cavity of 7-day-old SD rats by flushing of medium using a 22-gauge needle and cultured in complete growth medium (alpha-modified eagle’s medium (αMEM, GIBCO, USA), 10% fetal bovine serum (Sigma, USA), 100 U/mL penicillin and 100 mg/mL streptomycin) at 37 °C and 5.0% CO_2_ in a humidified incubator. The culture medium was replaced every 3 days. All experiments were performed with BMSCs at passage 3.

### mRNA Transfection in BMSCs 3D pellet

For 3D pellet culture of BMSCs, 1.5×10^6^ cells were aggregated into cell pellets by centrifugation at 1000 rpm for 5 minutes and the supernatant was carefully replaced with 10 mL of fresh medium. Chondrogenesis assays were induced by the complete chondrogenic medium, consisting of DMEM high-glucose, 10 ng/mL TGF-β3, 100 nM dexamethasone, 50 μg/mL ascorbic acid 2-phosphate, 1% ITS+ supplement, 10% FBS and 1% P/S. In mRNA treated groups, 10 ng/mL TGF-β3 protein was replacement with TGF-β3 mRNA. It was replaced with three-quarters of the medium every three days until day 21 in culture before taken for other assays.

### Efficiency and cytotoxicity detection of cmRNA complexes

We choose EGFP cmRNA as a reporter to demonstrate the transfection efficiency in HEK 293T cells and BMSCs. HEK 293T cells and BMSCs were seeded at 1×10^5^ cells per well in 12-well plate, left to recover for overnight then transfected by EGFP cmRNA at N/P=0, 4, 6, 8, 10, and 12 respectively. 24h after the transfection, cells were captured by fluorescence microscope for imaging, and the fluorescence indensity of each image was quantified by ImageJ to determine the appropriate N/P ratio of transfection. Subsequently, the cell cytotoxicity detection was carried out followed by the above processes using CCK-8 assay (Beyotime Biotechnology, China). Briefly, the cells were treated by CCK-8 reagent solution at 10:1 ratio per well. 1h after incubating in incubator, the absorbance was determined spectrophotometrically at 450 nm using a SpectraMax iD3 (Molecular Devices, USA).

### In vivo bioluminescence assays

Luciferase cmRNA only and Luciferase cmRNA/PEI compounds were injected into both leg joint of rats(n = 6 per group). The compounds were first assembled by mixing PEI and Luciferase cmRNA at N/P=8(Luciferase cmRNA: 20-200μg). Bioluminescence imaging of rats was proceeded using an IVIS Spectrum system (PerkinElmer, USA) at 24h and 96h after the injection. Firstly, rats were anesthetized with isoflurane, and the fur of inspection area was shaved to avoid the luminescent signal interference from the white fur. Rats were placed supine in the IVIS chamber. Superposition of bright field and dark field photos can intuitively show the location and intensity of specific bioluminescence signal in rats.

### DMMB assay for GAGs content evaluation

GAGs content was determined by 1,9-dimethylmethyleneblue (DMMB) dye-binding assay. Briefly, serially diluted samples were prepared and the DMMB solution was added. The absorbance was measured at 530 nm using the VMax Kinetic ELISA microplate reader (Molecular Devices, Inc., Sunnyvale, CA). GAGs content was normalized to DNA content in each specimen and presented as GAGs per cell.

### Histological evaluation of cell pellets

3D pellet samples were fixed in 4% paraformaldehyde (12h), frozen and sectioned prior to histological evaluation. H&E (hematoxylin-eosin), Safranin-O, Alcian Blue staining were performed strictly as previously described. For immunohistochemical analysis, 10μm slices were incubated for 30 minutes in blocking solution to prevent nonspecific binding and were then incubated with primary antibodies overnight at room temperature. Rabbit anti-human polyclonal antibodies against collagen type II and Aggrecan (Developmental Studies Hybridoma Bank, Iowa City, IA) were used in this study. A goat anti-mouse secondary antibody (Vector Laboratories Inc., Burlingame, CA) was used for detection. The reaction products were visualized by the Vectastain ABC kit and the DAM Peroxidase Substrate Kit (Vector laboratories Inc., Burlingame, CA), according to the manufacturers’ instructions. All negative controls were done using the same staining without using primary antibodies.

### Orthotopic model of cartilage defect in SD rats

The experimental animal use protocol for this project was approved by the Institutional Animal Care and Use Committee (IACUC), Shenzhen Institute of Advanced Technology, Chinese Academy of Sciences (SIAT-IACUC-200403-HCS-YY-A1097-01). Seventy-five 6-month-old male SD rats were purchased from Beijing Weitong Lihua Laboratory Animal Technology Co., Ltd, where they were housed and looked after in the experimental animal house. Two weeks later, they were anesthetized with 2% w/v pentobarbital sodium, the knee of SD rats was exposed and a chondral-only defect of 2 mm in diameter and 1.5 mm in depth was generated on the surface of patellar groove using electric drill (BOSCH, Garmany, GBM13). The rats were randomly divided into five groups (n=5 each) : (1) Sham group, (2) Control group (defect but without injection), (3) CB group (defect and inject a mixture of Collagen I and BMSCs), (4) CmR group (Defect and inject a mixture of Collagen I and cmRNA-TGFβ3), and CBmR group (defect and inject a mixture of Collagen I, cmRNA-TGFβ3 and BMSCs).

Collagen I (FibriCol®, Catalog:#5133, Advanced BioMatrix, USA) from bovine skin was prepared as previously described, and neutralized collagen I was diluted to 3 mg/ml using PBS, followed by BMSCs (10^7^ cells/mL) and PEI-cmRNA (cmRNA-TGFβ3: 200 ug/mL) were homogenized in the collagen solution by physical stirring at 4 °C. The preparation and storage of the complex were carried out at 4 °C and used within 1 hour. Next, collagen loaded with cmRNA and/or BMSCs was injected into the defective area. 20 μg of cmRNA encoding TGF-β3 and 1×10^6^ BMSCs were used for each cartilage defect. *In vivo*-jetPEI® (Polyplus-transfection® SA, Strasbourg, France) was used as a gene vector to deliver cmRNA-TGF-β3. According to the commercial instructions, PEI-cmRNA polyplexes at nitrogen to phosphate ratios (N/P) of 8 were formulated by mixing 50 μL of *in vivo*-jetPEI® solution to 100 µL 1.5mg/mL cmRNA encoding TGF-β3 for 15 min at room temperature. For in vivo testing, 20 μg of TGF-β3 cmRNA was used to prepare the polyplexes and then mixed into 100 µL collagen(PureCol^®^, Advanced BioMatrix, USA) for each rat joint, prior to implantation.

### Micro-computed tomography (µCT) analysis

Rat knee was analyzed in µCT scanner (NEMO micro-CT, PINGSENG Healthcare, China) at 2, 4, and 6 weeks post-surgery. µCT scanning was performed with focus over the hindlimbs with a 90 kVP tube voltage and 65 μA current. 3D rendering and processing of CT images were performed by using a commercial image processing software (Avatar; PINGSENG Healthcare, Shanghai, China).

### Histological analysis

SD rats were euthanized by injection with an overdose of pentobarbital sodium at 2, 4, and 6 weeks after surgery and the hindlimbs were cut off and then fixed in 4% paraformaldehyde, and subsequently decalcified with a 14% ethylenediaminetetraacetic acid (EDTA, SIGMA, USA) solution for four weeks, then embedded and sectioned (4 μm thickness) prior to histological staining. H&E, Safranin O-fast green staining and immunohistochemical staining for COL2A1 (1:1000, Abcam, USA) were performed according to instruction manual. The pictures were captured in a double-blind manner by an Inverted optical microscope (Nikon, Japan).

### Statistical analysis

Statistical comparisons were made using Student’s T test between two samples, and one-way analysis of variance (ANOVA) was used to compare the means among different groups. Tukey’s test was used in the post hoc multiple comparisons. All data are presented as the mean ± SD, and a “*p”* value of less than 0.05 was considered significant. “*ns*” stands for “not significant.”

## Results

### TGF-β3 cmRNA promotes matrix deposition of BMSCs in vitro

Figure 1 summarizes the study design. The chemically nucleotide-modified mRNA encoding gene reporters(mWassibi, Luciferase) and TGF-β3 were manufactured.A robustly of the mRNA expression both in vitro and in vivo was recorded(supplement Figure 1-3).One-shot injection of chemically nucleotide-modified Luciferase mRNA to the rat joint showed robust expression within joint and within 24 hours and could last at least 96 hours.Importantly there is no systematically exposure(Figure 2).Clearly,the chemically nucleotide-modified mRNA could be introduced in vivo at the right time and right place.

**Figure 1.**
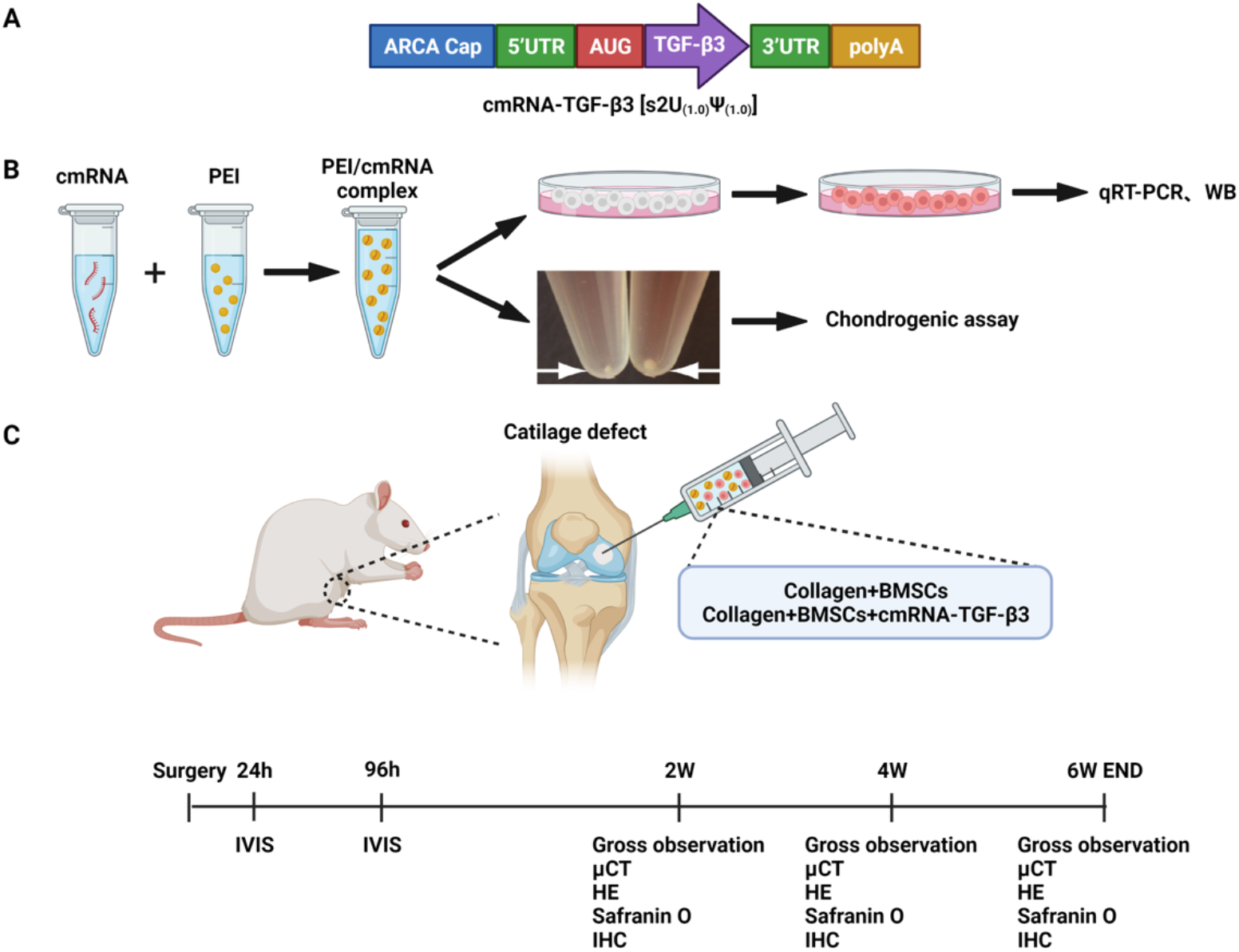
Experimental design. (A) Structure of TGF-β3cmRNA, (B)Evaluation of cmRNA both in monolayer culture and 3D micro mass culture of BMSCs, (C)Orthotopic model of cartilage defect in rats and the time line of experiments in vivo.

**Figure 2.**
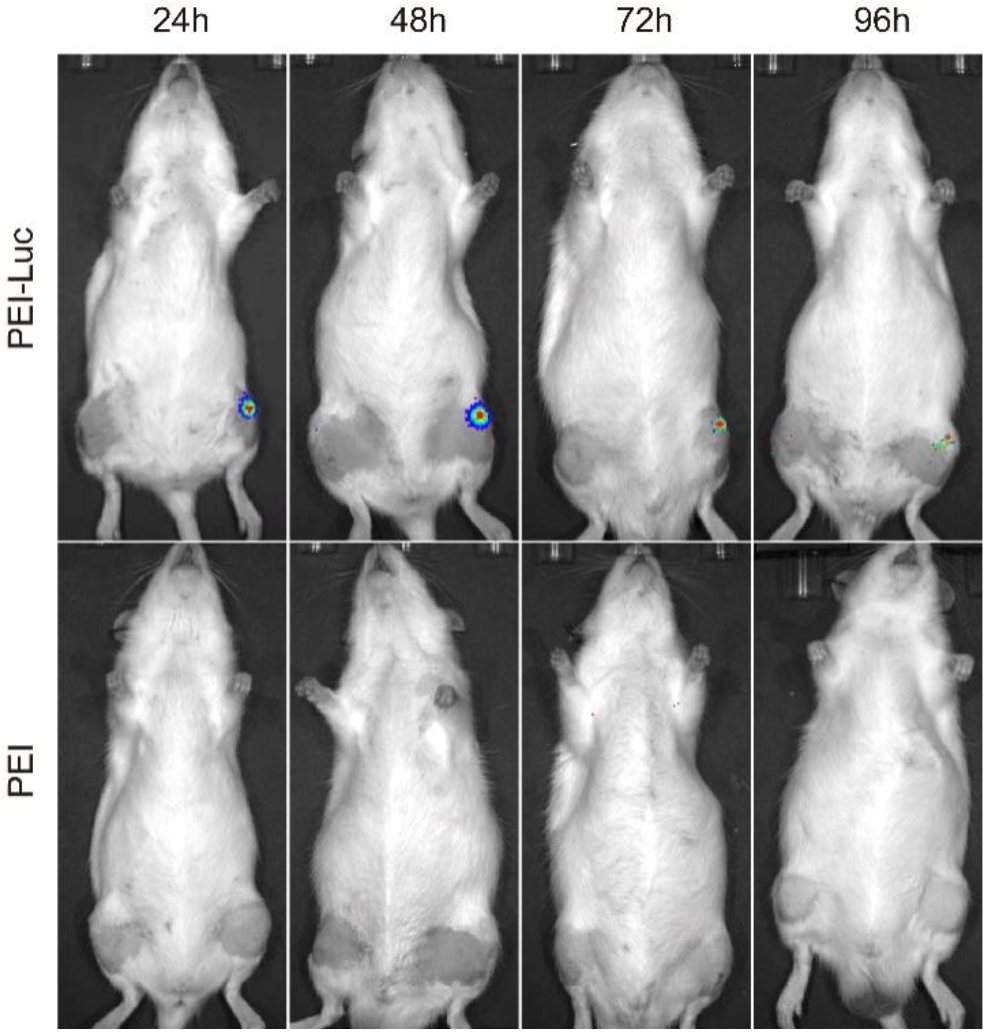
In vivo transfection and sustained epxression of cmRNA-Luciferase.

Then, we evaluated the ability of transfection and inducing chondrogenic differentiation of BMSCs in vitro. Besides, we used a standardized rat cartilage defect model to detect the therapeutic effect of TGF-β3 cmRNA in vivo. TGF-β3 plays an important role in the proliferation and matrix deposition of BMSCs. In addition to mono-layer cell culture, the transfection ability of TGF-β3 cmRNA on BMSCs was also verified in 3D culture system. As evidenced, strong signal of green fluorescence was observed and quantified in the spheroids of BMSCs treated with TGF-β3 cmRNA (Figure 3a, b). More importantly, the volume of pellet increased significantly in a time-dependent manner following TGF-β3 cmRNA treatment compared to vector group (Figure 3c, d). This indicated that TGF-β3 cmRNA also had excellent transfection ability in 3D culture system with the assistance of PEI vector, and the expression of TGF-β3 significantly enhanced the secretion of extracellular matrix of BMSCs.

**Figure 3.**
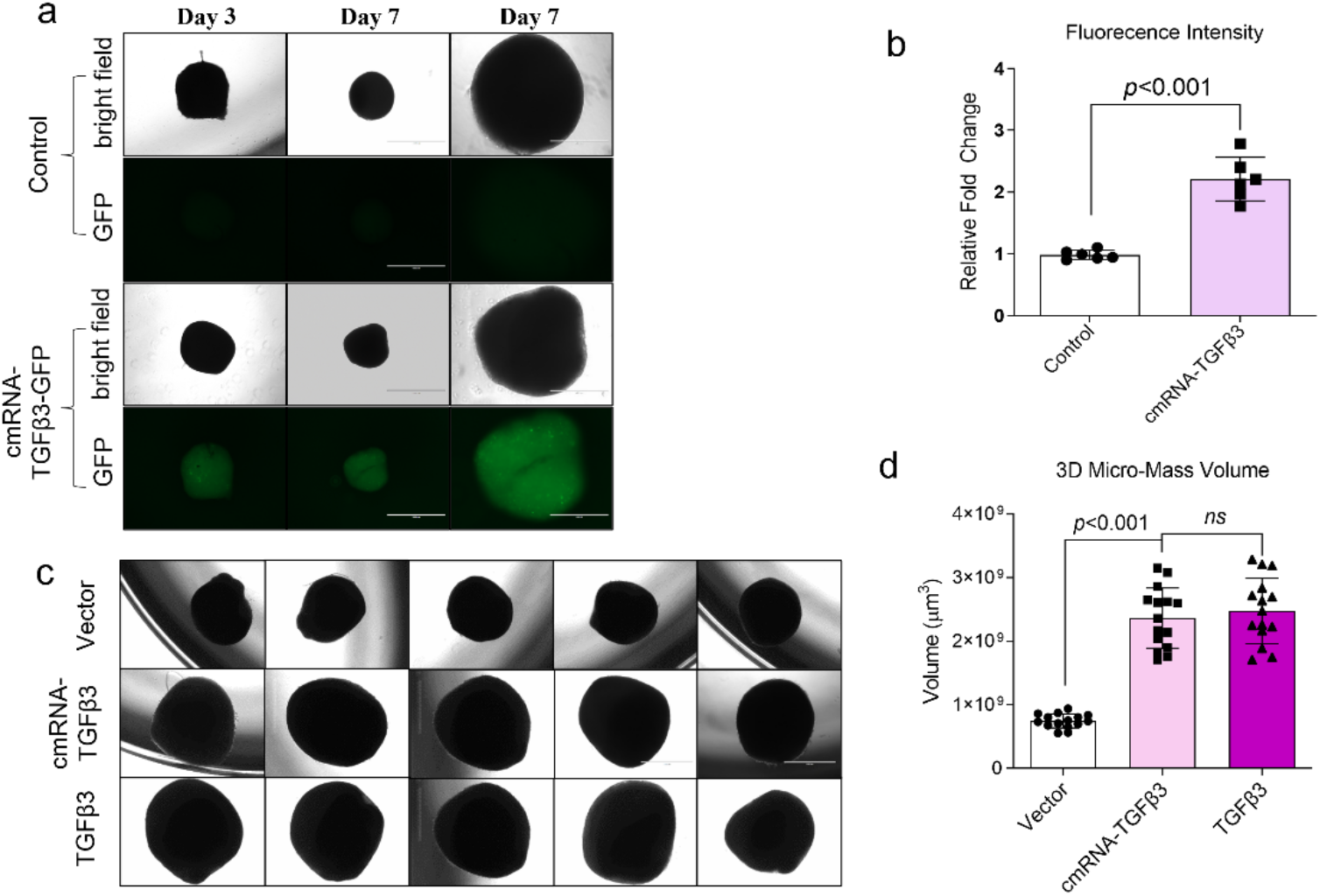
TGF-β3 cmRNA promotes matrix deposition for 3D micro-mass. Strong signal of green fluorescence was observed and quantified in the spheroids of BMSCs treated with TGF-β3 cmRNA (a, b). The volume of pellet increased significantly in a time-dependent manner following TGF-β3 cmRNA treatment compared to vector group (c, d).

### TGF-β3 cmRNA induces chondrogenesis of BMSCs in vitro

To examine the effect of TGF-β3 cmRNA on chondrogenic differentiation of BMSCs, histological staining was performed. Higher amount of proteoglycan deposition was observed in the TGF-β3 cmRNA treated samples, which displayed strong positive staining for Safranin O and Alcian blue staining. Additionally, abundant expression of cartilage-specific markers (collagen II and Aggrecan) showed that the chondrogenic differentiation of BMSCs was induced by TGF-β3 cmRNA successfully (Figure 4a). GAG are important components of cartilage ECM and are critical for bearing the load and reducing the friction cartilage experienced and endowing the lubricating properties of synovial fluid. Quantification of GAGs using DMMB assay showed that the TGF-β3 cmRNA transfected pellets yielded nearly 10-fold higher GAGs content than vector group and 2-fold higher than TGF-β3 protein treated pellet (Fig 4b). These results indicated that transfection of cmRNA-TGF-β3 induced the chondrogenic differentiation of BMSCs *in vitro*, even surpassed the chondroinductive effect of recombinant TGF-β3.

**Figure 4.**
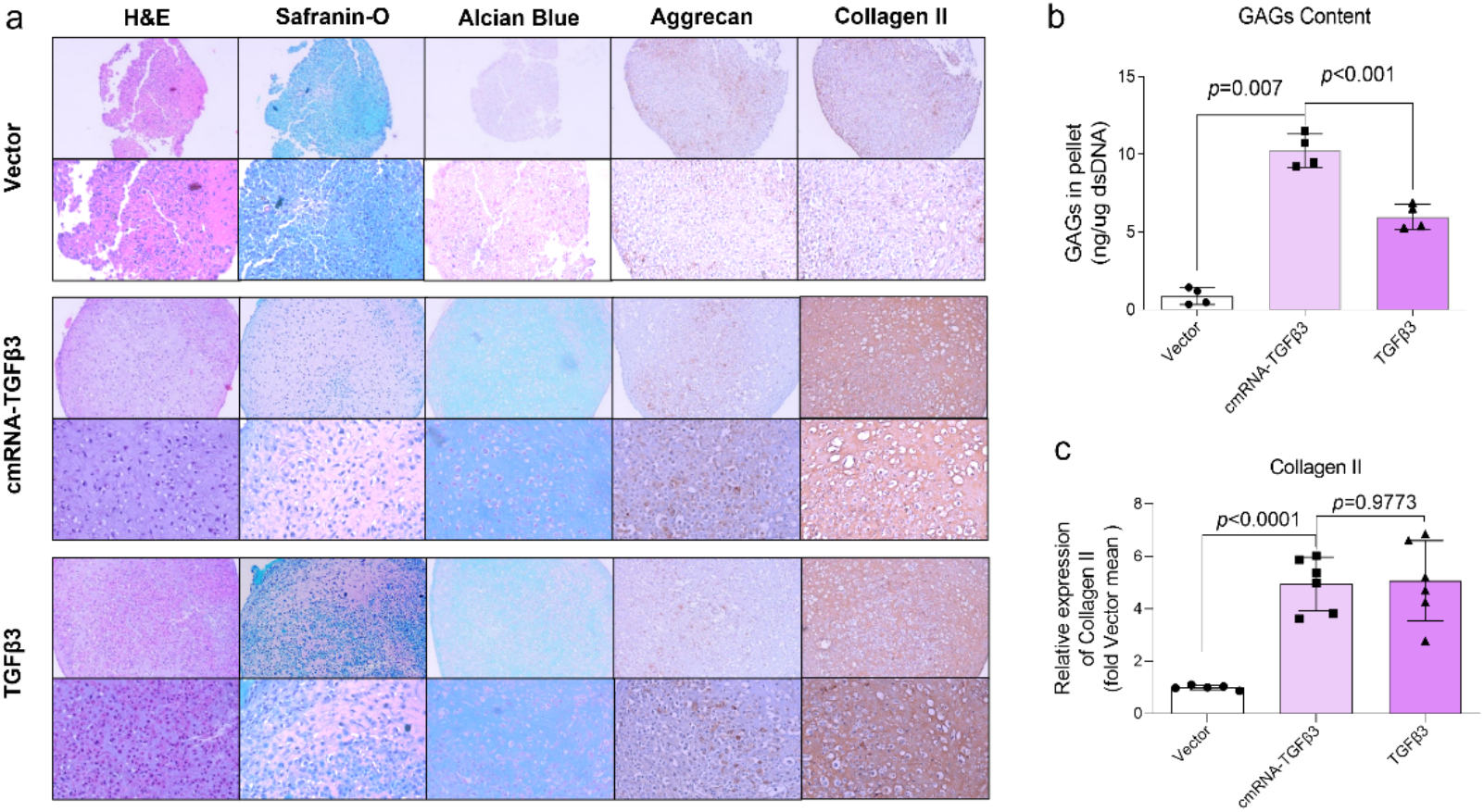
BMSCs pellets were treated 21 days later, (a)Histology and IHC Staining of Cartilage tissue(H&E, Safranin-O, Alcian Blue, Aggrecan, Collagen II) and GAGs content quantification. (b)GAGs content of BMSCs pellets.(c) Immunohistochemical (collagen II)semi-quantitative analysis.

### Macroscopic evaluation of TGF-β3 cmRNA promoting the repair of cartilage defect in rat model

After verified the integrity of TGF-β3 cmRNA, its good transfection efficiency in mono-layer and 3D pellet culture and chondroinductive properties for BMSCs in vitro, we next sought to test the ability of the TGF-β3 cmRNA to promote joint regeneration in a rat model of chondral defect repair. To this end, cylindrical chondral defects were created in the lateral trochlear ridge of hind limbs in 6-month-old SD rats. After creating a knee cartilage defect by open surgery, collagen loaded with cmRNA-TGF-β3 with or without BMSCs was injected into the defect site. Macroscopically, the control group showed low levels of self-repair, the cartilage defect was clearly visible two weeks after the operation, and there were large amounts of inflammatory exudation and rough joint surface can be observed. The CB group and the CBmR group demonstrated smoother joint surface and improved cartilage regeneration at the selected time point (Figure 5a). Four and six weeks after surgery, the control group showed some improvement compared to 2 weeks after surgery, however they displayed lower levels of defect fill higher tissue adhesion compared to CB group and the CBmR group at the same time point. We further used the International Cartilage Repair Society (ICRS) scale for macroscopic scoring of joint tissue, and this is in accordance with previous analysis (Figure 5b).

**Figure 5.**
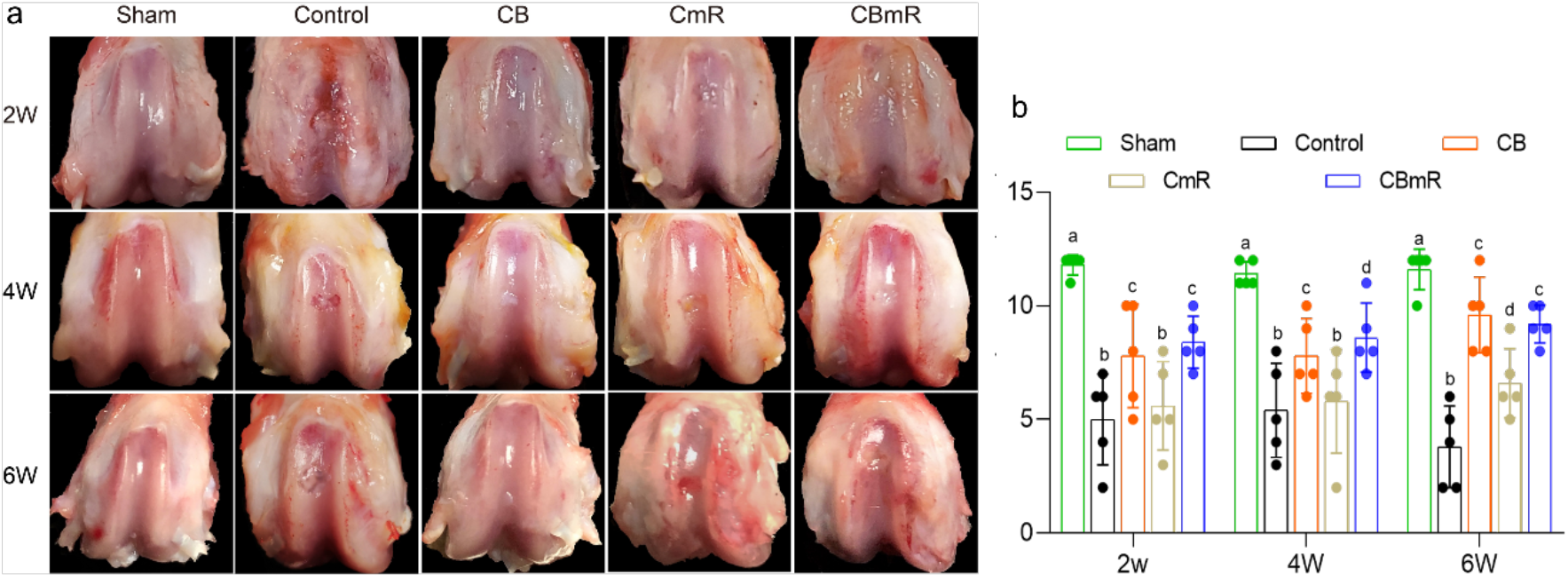
In vivo cartilage regeneration in defects with treating cmRNA-TGFβ3.(a) Gross images of femurs collected from different groups, The “Sham group” represented only exposure of the joint cavity, and no cartilage defect was created.; “Control group” stands for cartilage defect creation and injection of PBS: “CB group” stands for cartilage defect creation and inject mixture of Collagen I and BMSCs: “CmR group” stands for cartilage defect creation and inject a mixture of Collagen I and cmRNA-TGFβ3: “CBmR group” stands for cartilage defect creation and inject mixture of Collagen I, cmRNA-TGFβ3 and BMSCs. (b) ICRS macroscopic scores (mean ± s.d., n = 5, bars with different letters are significantly different from each other at p < 0.05).

### Histological analysis of cartilage repair via cmRNA-TGFβ3

Histological evaluation using hematoxylin and eosin (H&E), safranin fast green and immunohistochemistry (IHC) staining of type II collagen were performed to demonstrate the histological changes of cartilage after 6 weeks of cartilage tissue treatment. As shown in Figure 5a, the control group exhibited obvious subchondral bone abnormalities, with significant loss of cartilage layer. CB group and CBmR group had obvious improvement compared with control group, but subchondral bone abnormalities were still clearly visible in CB group. In comparison, in the CBmR group, more Safranin-O positive matrix deposition could be observed in cartilage defect site with smooth articular surface, and decreased subchondral bone abnormalities. More importantly, after TGF-β3 cmRNA treatment, the nascent tissue deposited increased amount of type II collagen (Figure 6a, e), which was the predominate collagen in native articular cartilage, indicative of functional repair of cartilage damage. Further quantitative analysis (Figure 5b-e) also consistent with the above observation. The ICRS histological score showed that CBmR was significantly higher than the other two groups. The same conclusions were drawn in the filling of the defect and the evaluation of cartilage thickness. Taken together, compared to CB group, CBmR group better promoted cartilage regeneration and recapitulate the specific collagen composition of native cartilage.

**Figure 6.**
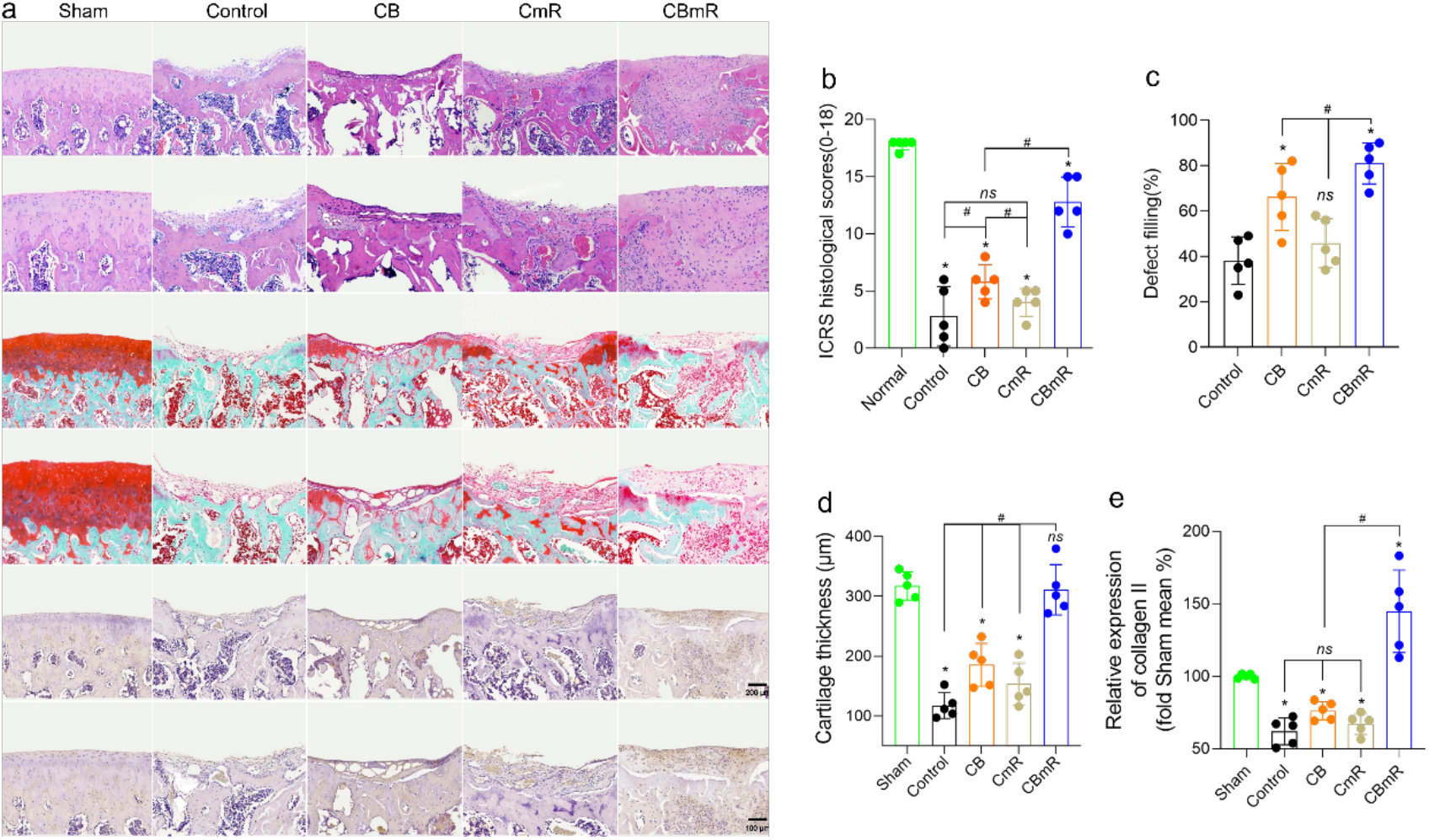
Histological analysis of cartilage repair by cmRNA-TGFβ3.(a) Histological analysis of OA was evaluated by HE, safranin fast green and immunohistochemistry staining.(b) ICRS histological score of articular cartilage was determined.(c) Defect filling (%).(d) cartilage thickness(μm).(d) Semi-quantitative analysis of immunohistochemistry (collagen II)by image J. Values are presented as means ± SD, n = 5 in each group. *P <0 .05, significantly different from control; #P <0 .05, significantly different; ns. not significant, as illustrated.

### Effects of cmRNA-TGF-β3 on bone changes in cartilage defects

When cartilage is damaged, it will lead to a series of degenerative changes in the subchondral bone. To examine the effect of cmRNA-TGF-β3 on subchondral bone during cartilage injury, microCT was performed. As shown in Figure 7, in the control group, the subchondral bone under corresponding cartilage defect site collapsed significantly. In addition, observed from the coronal section, the trabecular bone of the subchondral bone was thinned and lost. In the CB group and the CBmR group, the subchondral bone loss was significantly decreased, especially in the CBmR group, trabecular bone volume and bone mineral density is higher. In daily life, individuals encounter cartilage damage and are usually accompanied by joint involvement of subchondral bone. The subcondral abnormalities and bone loss would accelerate the occurrence of osteoarthritis and eventually lead to disability. Our results indicate that TGF-β3 cmRNA not only promoted cartilage regeneration but also inhibited the pathological changes of subchondral bone.

**Figure 7.**
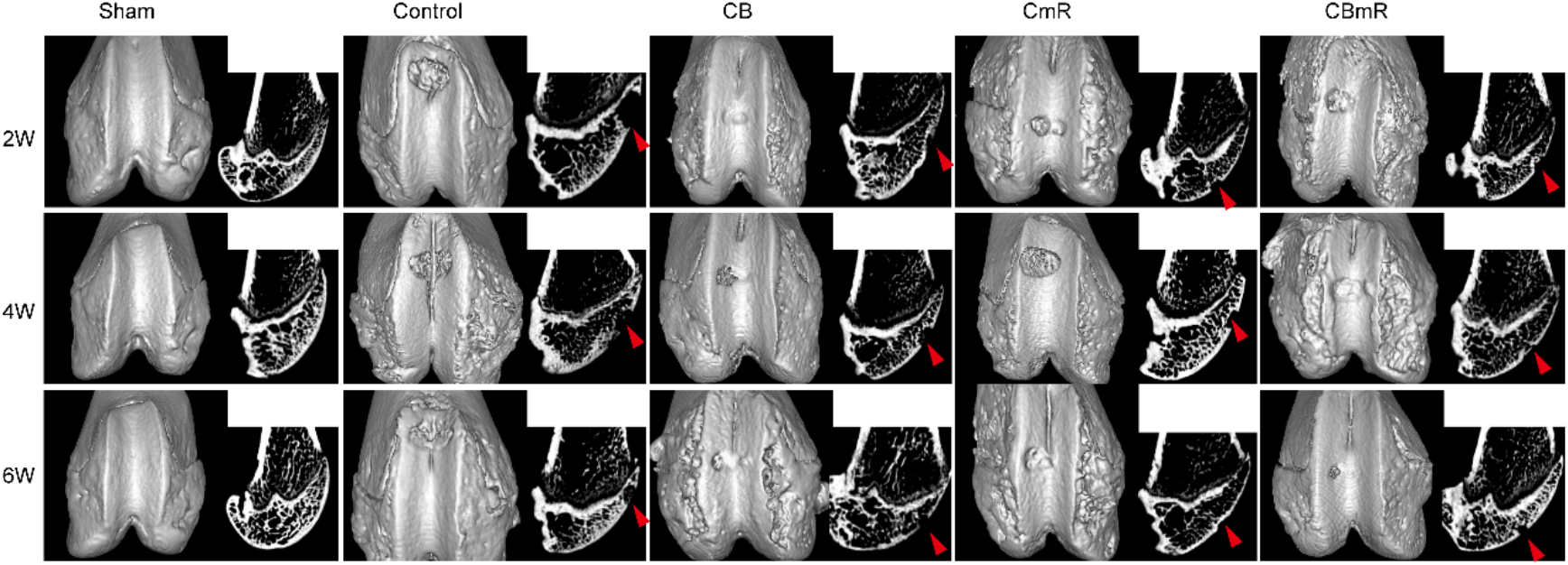
Micro-CT 3D reconstruction and longitudinal sections obtained from cartilage defect repair samples 2, 4, and 6 weeks post-treatment with cmRNA-TGFβ3 and/or BMSCs.

## Conclusion and Discussion

Our study advances an efficient approach to non-vascularized articular cartilage tissue rengeneration by introduction of pro-regenerative factors(s) in situ to enhance BMSCs therapeutic benefits. We demonstrate that chemically nucleoside-modified mRNA enchanced cell therapy is an effective, safe, and affordable approach to for potential clinical translation.

The articular cartilage is an indispensable part of the joint and is vulnerable to wear and tear or physical trauma such as sports or military associated injuries. Unfortunately, research has shown that damage to the cartilage can not self-heal or heal independently, due to the tissue’s low metabolic activity and avascularity. Without management, the damaged cartilage underwent progressive deterioration and posed negative effects on adjacent healthy tissue, ultimately landing to osteoarthritis (OA). Therefore there is neither cures nor disease-modifying manage for OA so far.

Facing the challegnes of stem cell therapies regarding cartilage tissue regeneration, the application of growth factors offered a straight forward solution, yet their successful clinical translation was hindered by disadvantages from many aspects. Therefore, we proposed a effective alternative by using chemically nucleoside-modified mRNA encoding a chondrogenic growth factor, TGF-β3, and packaged it into bone marrow mesenchymal stem cells(BMSCs) for in situ articular cartilage regeneration. TGF-β3 is an well-studies molecular target for guiding stem cells and chondrocytes differentiation, which has the ability to promote proteoglycan synthesis and to prevent hypertrophic chondrocyte differentiation and to conteract the negative effects posed on cartilage matrix and other adjacent tissue of IL1-β and other catalytic factors within the joint.

In this proof-of-concept study, after single injection of BMSCs bearing the modified TGF-β3 mRNA into the critical-sized cartilage defects of rats, an near complete healing of articular cartilage was achieved within 4-6 weeks (Figure5-7), with a better outcome compared with BMSCs only treatment group. Particularly, cartilage specific histological staining illustrated extracellular matrix deposition and integration of the regenerated tissue with the normal surrounding cartilage 4-6 weeks after the injection of the chemically nucleoside-modified mRNA enhanced cell therapy. In addition to cartilage regeneration, this TGF-β3 mRNA enhanced BMSCs therapy could prevent the joint trauma from progressing to subchondral bone damagnes which is a hallmark of late stage OA (Figure 7). Our findings suggested strongly this approach may be disease modifying measurement for OA. These findings proved, for the firt time that the BMSCs therapy can be enhanced dramatically by modified mRNA and the damaged avascular cartilage tissue can be restored in situ without systematical exposure of the modified mRNA, which may hold strong promise as a therapeutic platform for regeneration of other tissue.

Although compelling, this study does have some limitations. The cartilage defect was established in young, healthy rats which could fit certain types of trauma patients but not other type of patients such as elderlhy individuals which are major suffers of OA. In addition, whether the regenerated cartilage bear the biomechanical properties comparable to its nature counterparts also needs futher investigation performed in a relatively longer and larger study cohort. Lastly, the heterogeneity of BMSCs also needs to be considered while used side by side with cmRNA. Before planning clinical translation of our strategy, similar studies need to be carried out in a large animal models like porcine for further confirmation of its efficacy, safety and pharmacokinetics.

With two modified mRNA-based COVID-19 vaccines currently approved for clinical use along with a big success [27, 28], the safety and scalability of this technology has been established and modified mRNA technology has been applied in the treatment of various other diseases [29]. Tissue regeneration is a particularly suitable application for mRNA theraputics, because repetitive or systemic administration could be waived and the sustained regeneration of tissues could be achieved by effcient local growth factor expression in a timely and regional manner as illustrated by successful cases in liver, heart, skin and bone tissue [30-32] some of which has already advanced into clinical trials. These studies proved that mRNA can be used at a lower dose than its recombinant protein analog while obtaining successful therapeutic effects. Although systematical administration of nucleosides-modified mRNA could cause severe complications such as liver injury [33, 34] or heart problems[35]. Local injection in the context of regenerative medicine took advantages while avoided disadvantages of mRNA theraputics which clearly addressed their potential for successfully clinical translation.

Furthermore, the clinical application of mRNA have to overcome two main hurdles :the poor pharmacokentics (unstable) and high immunogenicity. Several methods were designed to resolve the problems such as modifying the 5′ cap, the 5′ and 3′UTRs, and the poly(A) tail[36, 37]. We previously reported the combination of the special UTRs and pseudouridine based modified for high translational efficiency and lower immunological responses to mRNA[38]. In this study, we provided proof-of-principle evidence for the first time that administration of chemically nucleoside-modified mRNA encoding TGF-β3 enhanced BMSCs into rat joints is capable of healing critical-sized defects of articular cartilage in situ through a superior way to its recombinant protein counterpart without systematically exposure.This technology could consolidate affordability, efficacy, safety with clinical expediency and break the barriers of DNA or recombination protein based gene therapy or systematically administrated mRNA therapy to clinical translation.

With futher development of mRNA therapeutics and its delivery technologies, we believe that clinical translation of mRNA therapeutics will be accelerated by the its improved effective, safe, not only in vaccine development or tunor therapeutics, but can also play an important role in regenerative medicine either alone or in combination to enhance cell therapies.

## Competing Interest Statement

YY received the research funding from:National Nature Science Foundation of China (82002355), Shenzhen Institute of Synthetic Biology Scientific Research Program (Grant No. DWKF20190010, JCHZ20200005), FC received the research funding from: the Shenzhen Institutes of Advanced Technology Innovation Program for Excellent Young Researchers (Y9G075), GeneHeal Medicine-SIAT mRNA Regenerative Medicine Laboratory (E1Z124). LZ received the research funding from: the National Natural Science Foundation of China [81660364, 81760343], Jiangxi Provincial Department of Education Research Program Major Project (171352), Science and Technology Planning Project of Jiangxi Health Commission (20191025, 202130131). XZ received the research funding National Nature Science Foundation of China (32071341). GW, JH, FY are employee at Gene Heal Medicine, a privately owned company developing mRNA therapeutics.All others have no potential conflict of interest.

## Acknowledgements

The authors thank the study participants and the authors thank Yuting Yang, Yuanyuan Zhang and Shi Chen for administrative support.

## Supplemental information

### Method and Results

#### Preparation of chemically modified mRNA(cmRNA) encoding TGF-β3, luciferase, TGF-β3-Wasabi

All oligos were synthesized by Integrated DNA Technologies (IDT, Coralville, IA).To generate templates for in vitro transcription, TGF *-β3* ORF DNA was synthesized by IDT.The Luciferase or wasabi ORF DNA purchased from Allele Biology(San Diego, CA) and subclone with UTR sequence to pcDNA 3.3 plasimid. Splint-mediated ligations were carried out using Ampligase Thermostable DNA Ligase (Epicenter Biotechnologies, Madison, WI). UTR ligations were conducted in the presence of 200 nM UTR oligos and 100 nM splint oligos, using 5 cycles of the following annealing profile: 95°C for 10 seconds; 45°C for 1 minute; 50°C for 1 minute; 55°C for 1 minute; 60°C for 1 minute. A phosphorylated forward primer was employed in the ORF PCRs to facilitate ligation of the top strand to the 5′ UTR fragment(5′UTR sequence: AAATAAGAGAGAAAAGAAGAGTAAGAAGAAATATAAGAGCCACC). The 3′ UTR fragment(3′ UTR sequence:GCTGCCTTCTGCGGGGCTTGCCTTCTGGCCATGCCCTTCTTCTCTCCCTTGCACCTGTACCTCTTGGTCTTTGAATAAAGCCTGAGTAGGAAGT) was also 5′-phosphorylated using polynucleotide kinase(New England Biolabs, Ipswich, MA). All intermediate PCR and ligation products were purified using QIAquick spin columns (Qiagen, Valencia, CA) before further processing. Template PCR amplicons were sub-cloned using the pcDNA 3.3-TOPO TA cloning kit (Invitrogen, Carlsbad, CA). Plasmid inserts were excised by restriction digest and recovered with SizeSelect gels (Invitrogen) before being used to template tail PCRs(Forward primer: TTGGACCCTCGTACAGAAGCTAATACG; Reverse primer: TTTTTTTTTTTTTTTTTTTTTTTTTTTTTTTTTTTTTTTTTTTTTTTTTTTTTTTTTTTTTTTTTTTTTTTTTTTTTTTTTTTTTTTTTTTTTTTTTTTTTTTTTTTTTTTTTTTTTTTTCTTCCTACTCAGGCTTTATT CAAAGACCA). Using MEGAScript T7 Transcription Kits (Life Technologies, Madison, Wis.) mRNA of TGF was synthesized and capped with the anti-reverse cap analog (ARCA:7-methyl (3’-O-methyl) GpppGm7G (5’)ppp(5’)G,New England Biolabs). To achieve mRNA modification, the following modified ribonucleic acid triphosphates were added to the reaction at a ratio of 100%: pseudouridine-5’-triphosphate and 5-methylcytidine-5’-triphosphate. Synthesized mRNA was purified and analyzed for size and purity.

#### Evaluation of cmRNA immunogenetic properties

After the TGFβ cmRNA was synthesized, the degree of immune response to unmodified and modified TGF-β3 mRNA were evaluated by transfection of the human Peripheral blood mononuclear cell, PBMC). RNA transfections were conducted using RNAiMAX (Invitrogen). RNA and reagent were first diluted in Opti-MEM basal media (Invitrogen). 100 ng/uL RNA was diluted 5x and 5 uL of RNAiMAX per microgram of RNA was diluted 10x, then these parts were pooled and incubated 15 minutes at room temperature (RT) before being added to culture media. Transfected cells were lysed using 400 uL CellsDirect reagents (Invitrogen), and 20 uL of each lysate was taken forward to a 50 uL reverse transcription reaction using the VILO cDNA synthesis kit (Invitrogen). Reactions were purified on QIAquick columns (Qiagen). qRT-PCR of the interferon related genes such as RIG-I (Forward primer: GTTGTCCCCATGCTGTTCTT;Reverse primer: GCAAGTCTTACATGGCAGCA) and TLR7 (Forward primer: CCTTGAGGCCAACAACATCT; Reverse primer: GTAGGGACGGCTGTGACATT). Transfected cells were lysed using 400 uL CellsDirect reagents (Invitrogen), and 20 uL of each lysate was taken forward to a 50 uL reverse transcription reaction using the VILO cDNA synthesis kit (Invitrogen). Reactions were purified on QIAquick columns (Qiagen). qRT-PCR reactions were performed using SYBR FAST qPCR supermix (KAPA) Biosystems).

#### Transfection efficiency of PEI-cmRNA polyplexes on HEK 293T cells

The transfection efficiency of PEI-cmRNA polyplexes was assessed by western blotting and RT-PCR. 1.0×10^5^ HEK 293T cells were seeded in 6-well plates, and after overnight, the medium was exchanged with serum-free medium. The polyplexes were assembled by mixing PEI and cmRNA-TGF-β3 at N/P=8 followed by an incubation for 15 min at room temperature. A complex containing 2 µg of TGF-β3 cmRNA was added to the med ium, and after 24 hours of treatment, cells were lysed by RIPA Reagent (Pierce, USA) containing 1% proteinase inhibitor (Thermo Fisher Scientific, USA) on ice for 30 min. The supernatants were harvested after centrifugation for 30 min at 12, 000 ×g. The bicinchoninic acid (BCA; Biocolors, China) method was used for estimation of total protein content. About 50 μL of protein samples were denatured and separated by 10% SDS-polyacrylamide gels and transferred onto nitrocellulose membranes (Millipore, USA). The membranes were blocked in 5% commercial skim milk at room temperature for 1 h and probed with primary antibodies for TGF-β3 (Abcam, USA, 1: 1000) or GAPDH (Abcam, USA, 1: 5000) at 4°C overnight, washed, and incubated with secondary antibodies (Cell Signaling Technology, USA, 1: 10000) at room temperature for 1 h. The membranes were visualized by enhanced chemiluminescence (ECL; Millipore, USA), and densitometry was performed using ImageJ software (Version 1.50i, USA).

#### Immunogenicity evaluation of cmRNA TGF-β3 with different chemical modification

For non-immunotherapy applications including protein replacement therapy investigated in this study, an important factor influencing the therapeutic outcome of mRNA is the inherent innate immune-stimulating activity, as it might promote mRNA degradation and inhibit the translation process. Thus, we first evaluated the influence of different chemical modification for TGF-β3 mRNA on the control of innate immune activation. Briefly, a linearized DNA template containing the TGF-β3 open reading frame, flanking 5′ and 3′ untranslated regions and a 100bp poly-A tail was constructed. ARCA capping and poly A tailing were performed to cater for stability and transfection efficiency of TGF-β3 mRNA. The immunogenicity of different chemically modified mRNA was detected. Exogenous single-stranded RNA arouses antiviral defenses in mammalian cells through interferon dependent pathway. As showed in Supplemental Figure 1a, complete substitution of either 5-methylcytidine (5mC) for cytidine, or pseudouridine for uridine in TGFβ3-encoding transcripts markedly attenuated interferon signaling as illustrated by qRT-PCR for the two of interferon related genes while the most significant enhancement was seen when both modifications were used together. And the half life of modified mRNA was dramatically prolonged compared to the unmodified one on the condition that the two modifications were done at the same time(Supplemental Figure 2b).

**Supplemental Figure 1.**
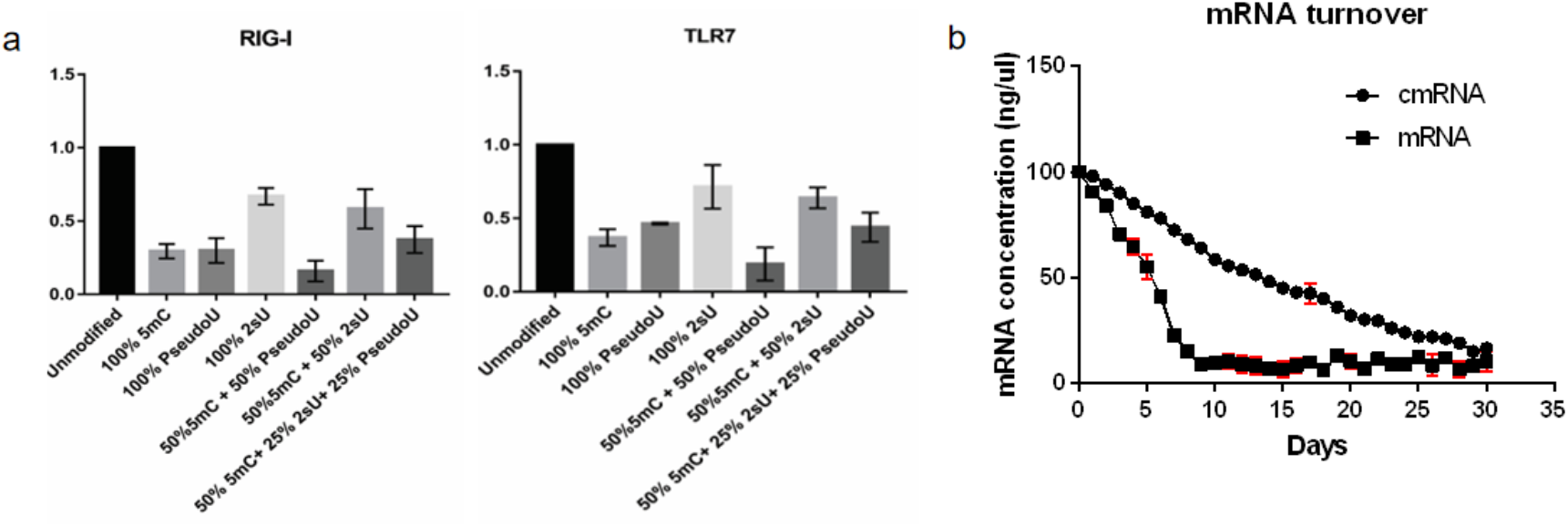
chemical modified mRNA overcomes antiviral responses in human PBMCs. (a)Quantitative RT-PCR data (normalized to unmodified mRNA) showing expression of 2 interferon-regulated genes in human PBMC after 24 hours transfection with unmodified and modified mRNA(pseudoU or 5mC at different ratio) encoding TGF-β3. (b)Half life of modified mRNA(pseudoU +5mC)in PCR tube without any treatment at 37°C.

#### PEI-cmRNA polyplexes efficiently deliver TGF-β3 cmRNA into HEK 293T cells

The expression kinetics of TGF-β3 cmRNA in HEK 293T cells were analyzed and compared at 24 h post-transfection using Western blotting and RT-PCR. As shown in Supplemental Figure 2 a,b, the expression of TGF-β3 was significantly increased. To further confirm the transfection efficiency of cmRNA in vitro, mWassabi reporter was utilized to label cmRNA, and a large amount of fluorescence signal was observed in cells treated with TGF-β3/mWassabi cmRNA (Supplemental Figure 2c). All the above results indicated that cmRNA had high transfection efficiency in monolayer cell culture system.

**Supplemental Figure 2.**
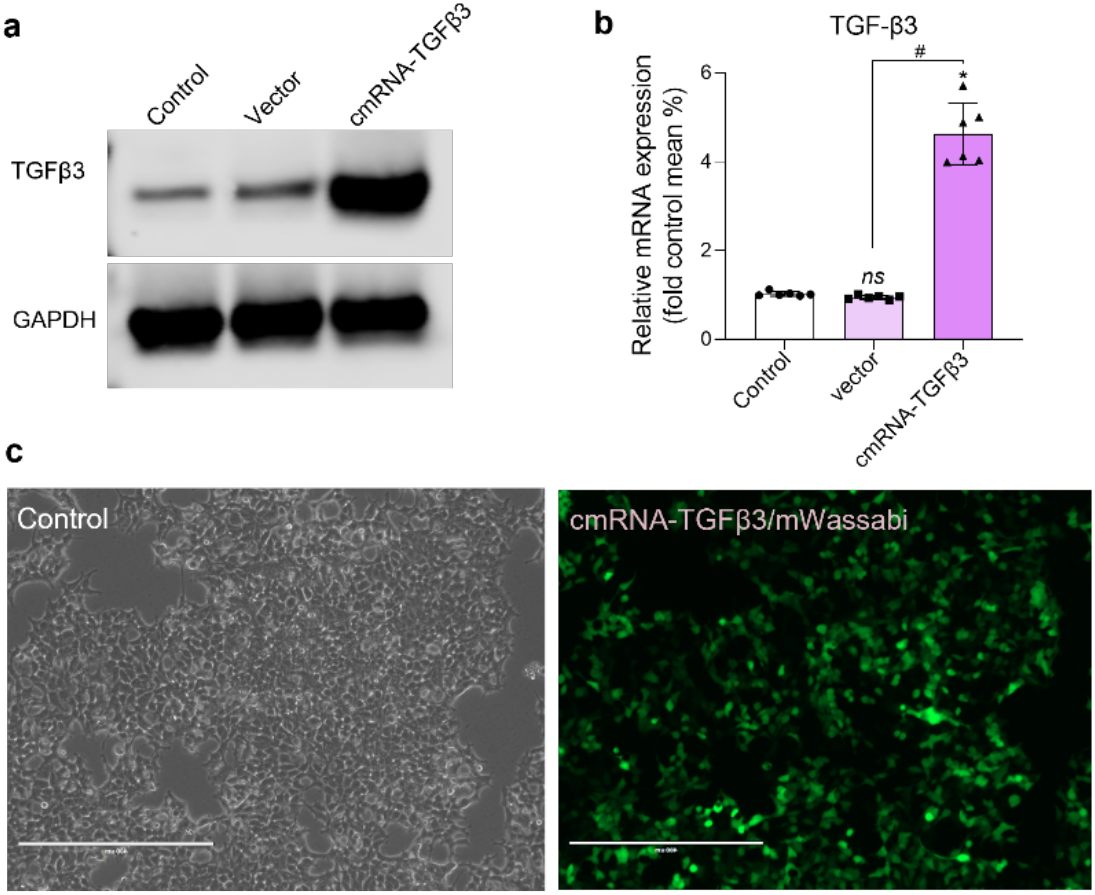
*In vitro* transfection efficiency of cmRNA-TGF-β3. HEK 293T cells were treated for 24 hours, (a) Western blot and (b) qRT-PCR analysis were used to detect the expression level of intracellular TGF-β3; (c) Wassabi was used to label TGF-β3, forming the PEI-cmRNA-TGF-β3/wassabi complex. Fluorescence traces were used to visualize the transfection ability of cmRNA-TGF-β3 after 24 hours of treatment of HEK 293T cells with the complex.

#### Efficiency and cytotoxicity detection of cmRNA complexes

To determine the appropriate N/P ratio of transfection, EGFP mRNA was transfected at N/P=4, 6, 8, 10, and 12 respectively in HEK293T cells and BMSCs. The fluorescence indensity showed that the efficiency of cell transfection was significantly improved when N/P=6 and 8 (Supplemental Figure 3a, b). Next, we evaluated the cytotoxicity of transfected HEK293T and BMSCs over 24h using a CCK-8 assay in order to confirm that the PEI-cmRNA complex has no negative effect on cellular growth. As shown in Fig. 4c, whatever the N/P ratio is, the proliferation of transfected cells was similar to the untransfected group, suggesting that PEI-cmRNA complex have no influence on cell proliferation.

#### Efficiency and cytotoxicity detection of cmRNA complexes

**Supplemental Figure 3.**
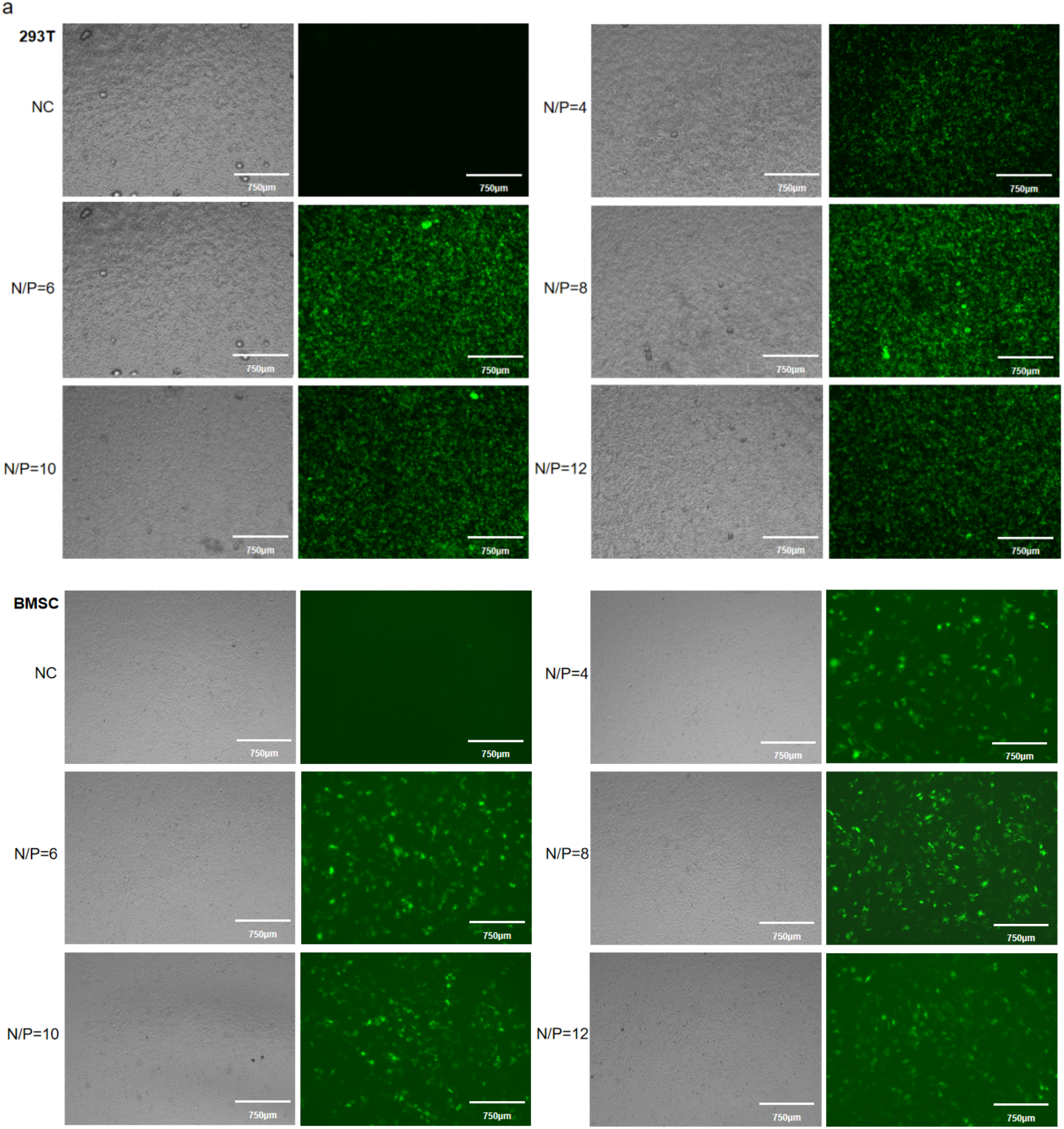
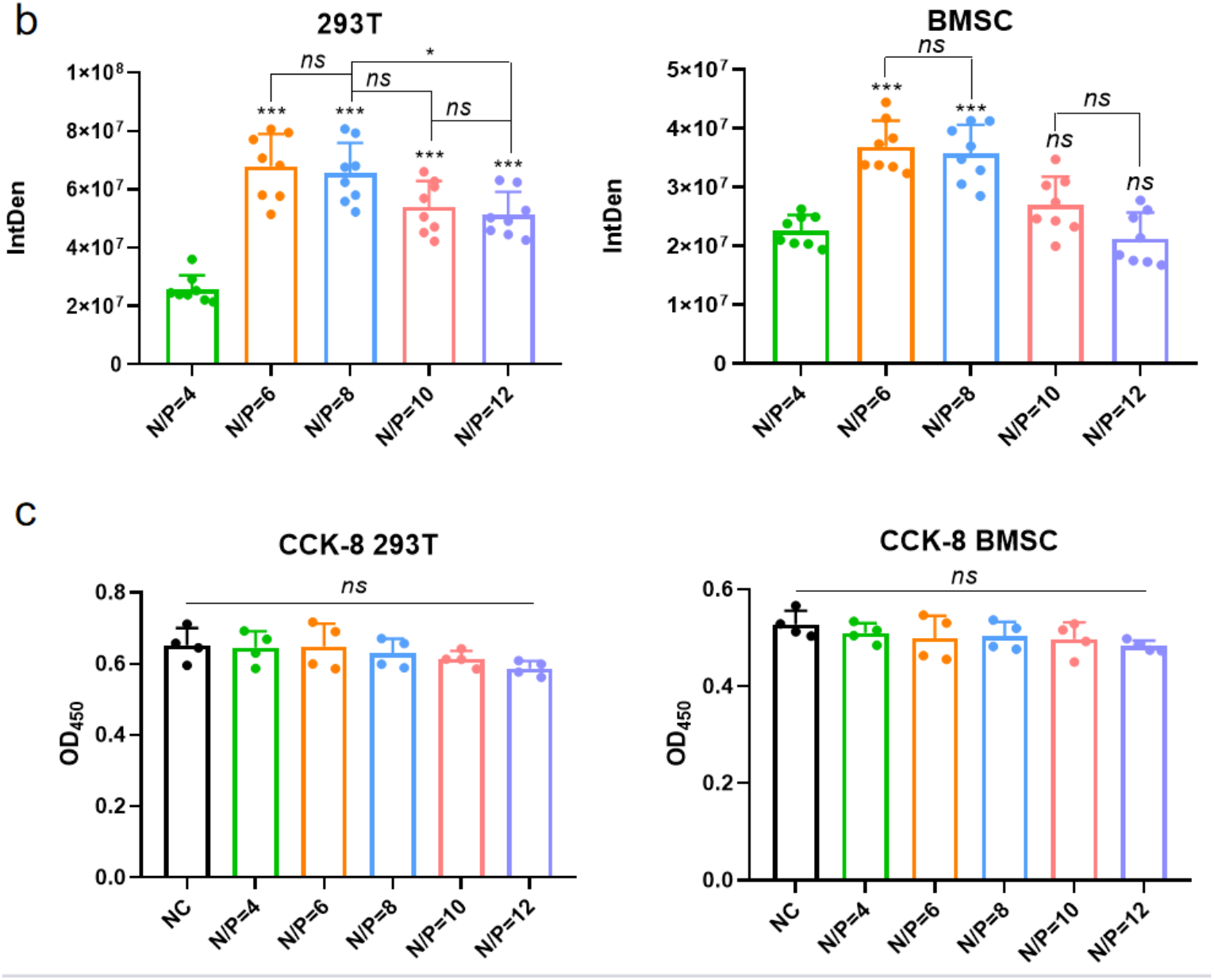
Efficiency and cytotoxicity detection of cmRNA complexes. (a) Representative fluorescence microscopy images of HEK293T cells and BMSCs taken 24 h after EGFP mRNA transfection at different N/P ration; (b) Fluorescence intensity of microscopy images was quantified by ImageJ; (c) Effect of transfection on HEK293T cells and BMSCs cytotoxicity.) Immunohistochemical (collagen II)semi-quantitative analysis.

